# Look4LTRs: A Long terminal repeat retrotransposon detection tool capable of cross species studies and discovering recently nested repeats

**DOI:** 10.1101/2023.07.28.551030

**Authors:** Anthony B. Garza, Emmanuelle Lerat, Hani Z. Girgis

**Affiliations:** Department of Electrical Engineering and Computer Science, Texas A&M University-Kingsville, Kingsville, Texas, United States; The Biometrics and Evolutionary Biology Laboratory, University Lyon 1, Lyon France

**Keywords:** long terminal repeat, transposable elements, LTR-retrotransposons, recently nested, bioinformatics, machine learning

## Abstract

Plant genomes include large numbers of transposable elements. One particular type of these elements is flanked by two Long Terminal Repeats (LTRs) and can translocate using RNA. Such elements are known as LTR-retrotransposons; they are the most abundant type of transposons in plant genomes. They have many important functions involving gene regulation and the rise of new genes and pseudo genes in response to severe stress. Additionally, LTR-retrotransposons have several applications in biotechnology. Due to the abundance and the importance of LTR-retrotransposons, multiple computational tools have been developed for their detection. However, none of these tools take advantages of the availability of related genomes; they process one chromosome at a time. Further, recently nested LTR-retrotransposons (multiple elements of the same family are inserted into each other) cannot be annotated accurately — or cannot be annotated at all — by the currently available tools. Motivated to overcome these two limitations, we built *Look4LTRs*, which can annotate LTR-retrotransposons in multiple related genomes simultaneously and discover recently nested elements. The methodology of *Look4LTRs* depends on techniques imported from the signal-processing field, graph algorithms, and machine learning with a minimal use of alignment algorithms. Four plant genomes were used in developing *Look4LTRs* and eight plant genomes for evaluating it in contrast to three related tools. *Look4LTRs* is the fastest while maintaining better or comparable F1 scores (the harmonic average of recall and precision) to those obtained by the other tools. Our results demonstrate the added benefit of annotating LTR-retrotransposons in multiple related genomes simultaneously and the ability to discover recently nested elements. Expert human manual examination of six elements — not included in the ground truth — revealed that three elements belong to known families and two elements are likely from new families. With respect to examining recently nested LTR-retrotransposons, three out of five were confirmed to be valid elements. *Look4LTRs* — with its speed, accuracy, and novel features — represents a true advancement in the annotation of LTR-retrotransposons, opening the door to many studies focused on understanding their functions in plants.

## Background

Transposable elements (TEs) are genetic elements capable of replicating and inserting themselves into new positions within a genome. TEs were discovered in the 1940’s by Barbara McClintock [1] in the maize genome. There was controversy surrounding their involvement in genetic processes as they were initially believed to be junk DNA with no function within a genome. However, recent studies have shown that TEs can play significant roles in a genome [2–7].

TEs are found in nearly all eukaryotes — both animals and plants — and prokaryotes. The human genome is composed of nearly half of TEs [7, 8], in comparison to the coding regions which only make up approximately 1–2% of the genome. For the wheat (*Triticum aestivum*) genome, 85% is made of repeats including TEs [9].

TEs are classified into two classes: Class I and Class II. Class I is composed of retrotransposons. Class II is composed of DNA transposons. These two classes are separated by their mechanism of transposition; retrotransposons use a “copy-and-paste” method where they move through reverse transcription, using an RNA intermediate, while DNA transposons use a “cutand-paste” method where they move through a DNA intermediate [2]. Recent alternate classifications have been proposed; most of these classifications depend on TE internal genes [10].

Retrotransposons can be further split into two categories: Long Terminal Repeat (LTR) retrotransposons and non-LTR-retrotransposons. LTR-retrotransposons are characterized by their LTRs which are two very similar sequences found at the 5’ and 3’ ends of a TE and a polypurine tract found in the region flanked by the two LTRs. It is important to note that these LTRs have the same orientation and are not complemented or reversed. A primer binding site is found on the opposite end of the internal part from the polypurine tract [10,11]. Non-LTR-retrotransposons are composed of Long Interspersed Nuclear Elements (LINEs) and Short Interspersed Nuclear Elements (SINEs).

DNA transposons can be grouped by the method of transposition and the presence of Terminal Inverted Repeats (TIRs) [10,12]. TIRs are two similar sequences except one is the reverse complement of the other. The first group of DNA transposons includes elements with TIRs — barring Crypton elements. Miniature Inverted-repeat Transposable Elements (MITEs) are one such type of DNA transposon. The second group has DNA transposons that use a different method of transposition from the cut-and-paste mechanism. This groups includes Helitrons, which excise only one strand of DNA, and Mavericks, whose method of transposition is still not fully understood but likely utilizes both strands for replication unlike Helitrons [10].

The translocation of a TE into a genomic location may be unsafe. For example, suppose a TE inserts itself into a gene. This could cause the gene to become non-functional; it could also lead to the gene producing an altered protein with detrimental effects. A nonfunctional gene as a result of an insertion can be seen with Hemophilia A, where a LINE element is inserted into the Factor VIII gene [13]. Many genetic diseases are caused by TE insertions [14] such as colon cancer and neurofibromatosis type 1 [15].

LTR-retrotransposons are the most abundant type of TEs in plants [16], which serves to dramatically increase the size of plant genomes [17]. They generally come from the two super-families Ty1/*Copia* and Ty3/*Gypsy* [10, 18] as well as from the *Bel-Pao* superfamily. Retroviruses are very similar in structure to the LTR-retrotransposon [10]. However, unlike other LTR-retrotransposons which can only translocate within the same cell, retroviruses can move through different cells due to the addition of an envelope protein [19].

LTR-retrotransposons carry enhancers and promoters that can also affect the expression of nearby genes [2, 18]. Different types of stress such as heat can cause activity in LTR-retrotransposons, resulting in genetic diversity and evolution [20]. According to the study in [21], LTR-retrotransposons inserted into the introns of a gene can cause suppression of that gene. There is evidence suggesting that LTR-retrotransposons can co-opt sequences from genes for their own purposes and vice versa [3,18]. Further, LTR-retrotransposons make up a significant portion of the repeats at the centromeres of plants [18, 22].

Tools capable of locating LTR-retrotransposons in a genome are therefore of great interest to researchers. Tools for locating TEs can be classified into six types: library-based, learning-based, signature-based, comparative-genomics-based, de-novo-based, and pipelines of tools [11, 23]. Detection of LTR-retrotransposons currently are only available through library-based, signature-based, and pipeline tools. Library-based tools such as RepeatMasker [24] use a database of known TEs such as Repbase [25] and Dfam [26] to identify LTR-retrotransposons. A known issue with library-based tools is that they are unable to identify novel LTR-retrotransposons. Signature-based tools such as LTRharvest [27, 28] and LtrDetector [29] identify TEs by their structural features, e.g., matching LTRs, polypurine tract, etc. Pipelines rely on multiple tools from the other categories to identify LTR-retrotransposons. Two examples of this are TransposonUltimate [30] and LTRpred [31]. However, pipelines are known to be computationally expensive and time-consuming as they run multiple tools as well as difficult to install.

LTR retriever [32] and LTRdigest [33] are postprocessing tools. These tools do not directly locate LTR-retrotransposons. Instead, they process the otuput of an LTR-retrotransposon detection tool such as LTRharvest. They are used to filter out false positives from the predictions of other tools and build LTR-retrotransposon libraries. Examples of such tools include LTR retriever [32] and LTRdigest [33].

At this time, *LTR-retrotransposon detection tools do not take full advantage of information within a genome as well as information across species*. Most of the tools for annotating LTR-retrotransposons are designed for locating single, complete elements; such tools are not specifically designed for locating recently nested elements. Usually, another tool is applied to locating recently nested LTR-retrotransposons after applying a tool for detecting single elements [34, 35]. However, not all information is carried from one tool to the next; further, some tools are now obsolete and unavailable [35]. Another limitation of the currently available tools for LTR-retrotransposons is that some of them rely heavily on alignment algorithms, resulting in expensive computations. To solve these issues, we developed *Look4LTRs*, a signature-based tool focused on the detection of LTR-retrotransposons genome-wide. *Look4LTRs* utilizes genome-wide information about the repetitiveness of a sequence [23]. Such information can also be collected across multiple closely related genomes. *Look4LTRs* can locate recently nested LTR-retrotransposons with multiple levels of nesting. Finally, *Look4LTRs* uses minimal local alignment and a machine-learning-based approach for calculating global, pairwise identity scores efficiently [36]. These innovations make *Look4LTRs* the state of the art in detecting LTR-retrotransposons computationally.

## Implementation

### Input & Output

*Look4LTRs* accepts FASTA-format files. We suggest that *Look4LTRs* is given an entire genome at minimum. Multiple related genomes can be processed simultaneously by *Look4LTRs* to perform cross-species studies. The tool outputs the positions of long terminal repeat (LTR) retrotransposons, the location of the polypurine tract, and the location of the target site duplications.

### Data

A module of *Look4LTRs* — the detector — is an instance of supervised machine learning, which requires the availability of labeled examples for training and testing. For this reason, we needed some genomes for training the tool and others for testing it. For training our tool, we utilized the genomes of the following four species:

- *Arabidopsis thaliana* (TAIR10: thale cress)
- *Oryza sativa japonica* (IRGSP-1.0: japonica rice)
- *Glycine max* (Glycine max v2.1: soybean)
- *Sorghum bicolor* (Sorghum biclor NCBIv3: great millet)

For testing our tool, we utilized:

- *Zea mays* (AGPv4: corn)
- *Solanum lycopersicum* (SL3.0: tomato)
- *Solanum tuberosum* (SolTub 3.0: potato)
- *Theobroma cacao* (Theobroma cacao 20110822: cacao tree)

For testing the cross-species feature of our tool, we used these four species from the rice genus:

- *Oryza glaberrima* (Oryza glaberrima V1: african rice)
- *Oryza sativa indica* (ASM465v1: indica rice)
- *Oryza longistaminata* (ASM980554v1: longstamen rice)
- *Oryza rufipogon* (OR W1943: wild rice)

For manual inspection of the results, we used *Hordeum vulgare* (MorexV3: barley). These genomes are plants with high TE content. *Arabidopsis thaliana* was specifically chosen for training because it is a model organism with a well-annotated genome.

We utilized RepeatMasker [24] and a program called One Code To Find Them All [37] to locate Long Terminal Repeat (LTR) retrotransposon for our ground truth dataset. RepeatMasker is first used for locating LTRs and the internal parts of retrotransposons, using the Repbase database [25]. We used the slow search parameter to increase the sensitivity of the search and provided the appropriate species name for each genome. The only exceptions to providing the species names were the species chosen for the cross-species evaluation (barring *Oryza sativa japonica* and *Oryza sativa indica*). These species were not specifically well annotated in the Repbase library. Therefore, we instead passed the genus *Oryza*. The outputs of RepeatMasker were then passed to One Code To Find Them All, which assembled LTRs and the internal parts into complete LTR-retrotransposons.

We filtered these LTR-retrotransposons according to three criteria: (i) the LTRs and the internal parts must be at least 200 base pairs (bp) long individually, (ii) the LTR-retrotransposon must have at least 80% identity with a consensus sequence, and (iii) the LTR-retrotransposon must have at least 80% coverage of a consensus sequence. When performing any of the following checks, we account for nested elements in our calculations, allowing confirmed nested elements to remain in our ground truth. For example, suppose we have an element whose internal part is 1000 bp long and another element whose total length (LTRs and internal part) is 2000 bp. If the latter is nested within the former, then the size of the former’s internal part becomes 3000 bp. Presume that the consensus sequence for the former element has an internal part of 900 bp. The former element would be unable to pass the 80% identity or coverage checks. However, by not considering the nested element in these calculations, the internal part of 1000 bp would not change to 3000 and would thus pass the checks. Each LTR and each internal part are checked individually for a minimum length of 200 bp. If any of these regions is less than 200 bp, the whole element is discarded. The LTRs and the internal part in between are then individually aligned to their consensus sequence using Nucleotide BLAST [38]. From BLAST’s alignments, we select the longest alignment with at least 80% identity score. Any elements that failed to meet the minimum 80%-identity criterion are dropped. When checking the coverage, we sum the alignment lengths of the LTRs and their corresponding internal parts and compare the total length to the consensus sequence length. If the coverage is less than 80%, the element is dropped. The remaining LTR-retrotransposons are considered as our golden standard.

### Semi-Synthetic Genome Generation

Annotations of LTR-retrotransposons in the training genomes are incomplete, i.e., we do not know the locations of all LTR-retrotransposons in a genome. When training *Look4LTRs* on the training genomes, these unknown elements would complicate our training process. To deal with this problem, we generated semi-synthetic genomes for each training genome. We followed the following four steps to generate a semi-synthetic genome: (i) find LTR-retrotransposons for a genome according to our ground truth, (ii) extract these LTR-retrotransposons from the genome, shuffle the remaining regions randomly to destroy all other elements and repeats, and (iv) reinsert the LTR-retrotransposons at their original positions.

### Method overview

*Look4LTRs* annotates LTR-retrotransposons in a genome or *a group of related genomes*. The tool consists of these five modules (Figure 1):

**Figure 1.**
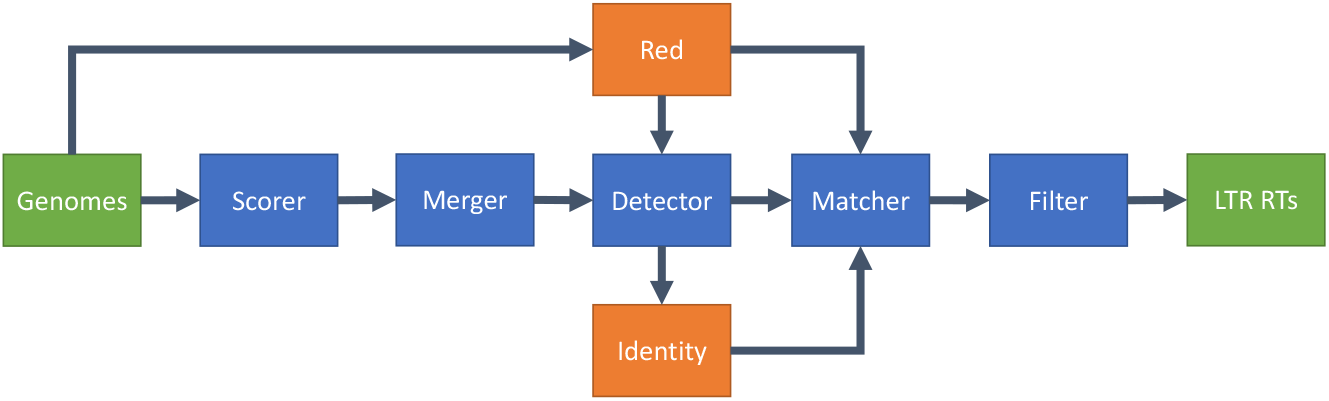
Overview of *Look4LTRs*. Our tool takes one genome or a group of related genomes as the input and outputs Long Terminal Repeat (LTR) retrotransposons. Five modules comprise *Look4LTRs*: (i) the scorer module, (ii) the merger module, (iii) the detector module, (iv) the matcher module, and (v) the filter module. The scorer module scores the genomes by nearby, matching k-mers. The merger module uses the scores to begin assembling potential LTRs. The detector module uses repeat content learned by *Red* (a tool for detecting all types of repeats uncategorized) from the genome(s) to finish assembling potential LTRs. The matcher module matches LTRs to each other to form LTR-retrotransposon candidates. The filter module removes candidates that fail to meet one of our confirmation criteria, e.g., the presence of a polypurine tract.

- **The scorer module** is responsible for scoring the genome by distances between k-mer copies,
- **The merger module** is responsible for merging similar regions of scores together into stretches,
- **The detector module** is responsible for merging stretches into candidate LTRs (not the entire LTR-retrotransposon) by a trained classifier,
- **The matcher module** is responsible for matching pairs of candidate LTRs into potential LTR-retrotransposons, and
- **The filter module** is responsible for filtering out potential elements that cannot be confirmed.

*Look4LTRs* output LTR-retrotransposons including *recently nested elements*.

The detector and the matcher modules utilize the tool *Red* [23], which can locate all types of repeats (tandem and interspersed) in a genome without grouping them into specific types or families. This allows *Look4LTRs* to use the repeat content found by *Red* to help connect stretches into candidate LTRs in the detector module. Additionally, *Look4LTRs* uses the repeat content for matching LTRs appropriately in the matcher module; specifically, if the internal part of an LTR-retrotransposon is not repetitive — that is to say, does not repeat throughout the genome — this candidate is unlikely to be an LTR-retrotransposon. The matcher module utilizes another tool called *Identity* [36], which calculates global identity scores of sequence pairs efficiently. These identity scores are incorporated as a part of our multi-evidence matching approach. *These two tools provide APIs, allowing for their integration with the other five modules into one code base*; *Red* and *Identity* are not called externally. The code of the five modules and the two APIs are packaged and shipped together. Thus, the user is not required to install multiple tools — just one. Next, we give the details of each module starting with the scorer.

### Scorer

The input to this module is a sequence of DNA and the outputs are two groups of scores, called the forward scores and the backward scores. The sequences are scored by 13-mer matching. This value of 13 was determined experimentally on our training genomes. We ran *Look4LTRs* on our training genomes with different values of k (10–15) and we found that 13 resulted in the best performance.

Our method of scoring was inspired by LtrDetector [29]. There, they score a k-mer by the distance to its closest match in either direction. However, this causes what we refer to as the *castle problem*. When there are multiple, sequentially inserted LTR-retrotransposons of the same family that are degenerate, it is expected for the k-mers of one LTR to have closest matches in any of the other LTRs, before or after it. This results in difficulty in determining if a region belongs to a single element. See Figure 2 for an example of the castle problem.

**Figure 2.**
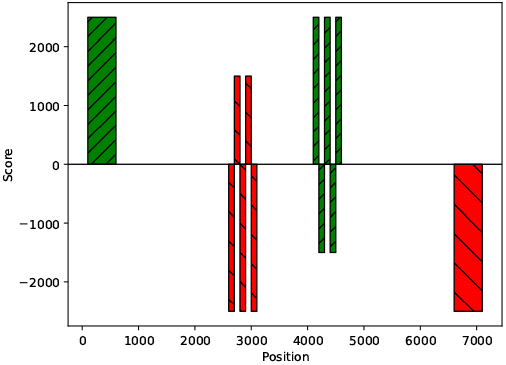
The castle problem. Represented are two LTR-retrotransposons of the same family. For simplicity, the internal parts were ignored. The 5’ LTRs are marked by diagonal hatching and the 3’ LTRs are represented by backward diagonal hatching. In this case, a singular vector of scores was used where positive y-values mark a distance to a matching k-mer in the forward direction and negative y-values mark a distance to a matching k-mer in the backward direction. Only the closest match is scored, regardless of direction. The 5’ LTR of the left element is clearly distinguished. The 3’ LTR of the left element is broken into positive and negative scores. The same holds for the 5’ LTR of the right element. The 3’ LTR of the right element is clearly distinguished.

To solve the castle problem, we depart from the method of LtrDetector and utilize a new scoring system. Where they have one score per k-mer, we have two scores per k-mer, one for the forward (downstream) direction and one for the backward (upstream) direction. In doing this, we minimize the castle problem, allowing for easier detection of LTRs later. This new scoring completely changes downstream analysis and should not be considered an incremental improvement. The forward scores are calculated by matching kmers forward, i.e., copies that are found downstream. The backward scores are calculated by matching kmers backward, i.e., copies that are found upstream. We first match every k-mer to its closest complete match (forward or backward), within a minimum distance of 400 and a maximum distance of 27,000. If a match is found within this range, we set the score of the current k-mer to the distance between itself and its copy. If no match is found, the score is set to 0. For example, suppose we have a k-mer starting at position 1,000 and a matching k-mer at position 2,000. Let’s refer to them as A and B. The score assigned to A in the forward scores is 1,000 as it is 1,000 bp away from B. The score assigned to B in the backward scores is also 1,000 as it is 1,000 bp away from A. For brevity, when talking about scores, we say that a k-mer is *pointing* to its copy. Note that each direction is scored separately. Next, we explain the rational of the scorer module. Consider a single LTR-retrotransposon. The 5’ LTR and 3’ LTR are similar to each other. That is to say, the k-mers in the 5’ LTR will have matches in the 3’ LTR and vice versa. In the forward scores, the 5’ LTR will have k-mers pointing to the 3’ LTR. In the backward scores, the 3’ LTR will have k-mers pointing to the 5’ LTR. Now, consider two recently nested elements, i.e., one element is nested inside another of the same family. In the forward and backward scores, the outer element will have k-mers pointing to the inner element and vice versa. However, the inner element will also have k-mers in its 5’ LTR pointing to its 3’ LTR. The forward and backward scores generated form a distinctive pattern that we later utilize to discover recently nested elements and single LTR-retrotransposons. See Figure 3 for examples of the forward and backward scores of LTR-retrotransposons.

**Figure 3.**
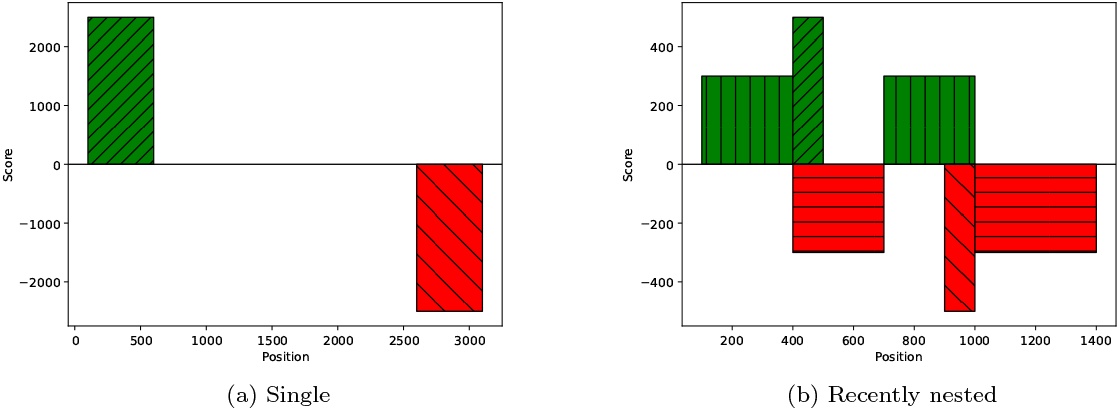
Forward and backward scores of LTR-retrotransposon(s). The y-axis shows the scores where positive values represent the forward scores and negative values represent the backward scores. The x-axis shows the genomic positions. (a) A single LTR-retrotransposon. The green bar is the 5’ LTR and the red bar is the 3’ LTR. The green bar has a score of approximately 2,600, meaning its match begins at approximately 2,600. (b) Two recently nested LTR-retrotransposons. The forward scores and backward scores of the nested LTRs are clearly distinguished. The 5’ LTR of the inner element is at positions 400–500 and the 3’ LTR is at positions 900–1,000. However, the forward and the backward scores of the LTRs of the outer element have merged with the internal parts. The outer 5’ LTR and a slice of the internal parts are at positions 0–400 and the outer 3’ LTR and a slice of the internal parts are at positions 1,000–1,400. Note that the scores of the outer element are pointing to the inner element. The outer 5’ LTR and a slice of the internal parts point to positions 400–700, which include the inner 5’ LTR and a slice of the inner LTR-retrotransposon’s internal part. The outer 3’ LTR and a slice of the internal part point to positions 700–1,000, which include the inner 3’ LTR and a slice of the inner LTR-retrotransposon’s internal parts. It is important to understand that in a recently nested LTR-retrotransposon, the outer LTRs do *not* point to each other but rather to the inner element.

Next we discuss how we merge our scores to form regions that may belong to LTRs.

### Merger

The forward scores and backward scores are passed to the merger module separately. It groups scores — forward or backward — into stretches; a stretch is a region of similar scores with possible small gaps, i.e., scores of zero. We refer to the median of non-zero scores in a stretch as that stretch’s height. We also refer to the number of bps in a stretch (or gap) as the size of the stretch (or gap).

To group scores into stretches, we performed statistical analysis of LTRs found in the training genomes. Two LTRs of the same element are unlikely to be identical because of mutations, which appear as gaps of zero scores. By calculating the distribution of the size of gaps within LTRs, we determined a value for the maximum difference between two scores to be considered similar, i.e., belong to the same LTR. We assumed the gap size is normally distributed. The mean size of a gap within an LTR was found to be 16.30 bp. The standard deviation was 19.62 bp. By adding 3 standard deviations to the mean (covering up to 99.7% of a normal distribution), we determined the similarity margin to be 75 bp. For two scores to be considered similar, the absolute difference between them must be less than the similarity margin.

The merger module performs the following steps:

1. Group regions of the same exact non-zero scores into stretches.
2. Label these stretches by size; stretches with a size of 16 or more are labeled as keep stretches, otherwise they are labeled as delete stretches. This method of labeling was inspired by LtrDetector [29].
3. Merge stretches with similar scores (the absolute difference in height is less than 75) provided that the gap between them is less than 75 (the similarity margin).
4. Remove interruptive stretches, which are delete-labeled stretches in between two stretches that would be merged if an interruptive stretch was not in between them.
5. Merge stretches.
6. Remove delete stretches.
7. Merge stretches.

As mentioned before, this merging process is applied to both the forward scores and the backward scores independent of one another. The forward scores become the forward stretches, each one pointing to a location further downstream. The backward scores become the backward stretches, each one pointing to a location further upstream.

### *Red* Training

*Red* is a self-supervised, hidden-Markov-model-based tool that can detect repeats (interspersed and simple) in a genome. It does not group repeats — including LTR-retrotransposons — into families. *Red* outputs a score for each k-mer in the input genome. This score indicates how many times above what is expected by chance a specific k-mer occurs in a genome. *Red* gives a value of zero to a k-mer that occurs less than what is expected by chance. We refer to the score as *Red* score. With respect to software integration, *Red* is accessed through an API and is *not* called externally. The code for *Red* is integrated into *Look4LTRs*.

### Detector

The detector module is responsible for the final step in the merging stage. It takes the forward stretches and backward stretches independently. The detector merges stretches into LTR candidates.

A linear classifier — trained by the stochastic gradient descent algorithm — is utilized to predict whether consecutive stretches should be merged. Each pair of consecutive stretches is described by the following 10 features:

- Size of the first stretch.
- Size of the second stretch.
- Size of the gap between the stretches.
- Absolute difference in height between the stretches.
- The absolute difference between the *Red* score medians (not counting zeros) of the two stretches.
- The absolute difference between the *Red* score means of the two stretches.
- Mean *Red* score of the first stretch.
- Mean *Red* score of the second stretch.
- Mean *Red* score of the gap between the stretches.
- Whether both stretches lie within the same repetitive region predicted by *Red*.

We split our data into three sets: training, validation, and testing, consisting of 70%, 20%, and 10% of the stretches found in the four training genomes. We subtracted the mean and divided by the standard deviation of each feature (except the last feature because it is binary). The mean and the standard deviation were calculated on the training set. To determine the parameters of the model, a random search 10-fold cross validation was performed on 1,000 iterations. We trained the classifier with the parameters found by the random search and kept the model with the best F1 score on the validation set. We performed a final evaluation on the testing set. After determining that our model could achieve satisfactory results, we rescaled the data and trained the classifier on the entire dataset. The final classifier achieved a recall of 92.81%, a precision of 64.37%, and an F1 score of 76.02%.

The trained classifier is given consecutive pairs of stretches and determines whether each pair should be merged. Any stretch that fails to merge with others is still considered an LTR candidate, which consists of one stretch. Candidates shorter than 200 bp (the minimum size of an LTR) are discarded.

### *Identity* Training

*Identity* is a machine-learning-based tool designed for predicting pair-wise global identity scores efficiently [36]. *Identity* takes a database of the sequences that will be compared later. Because *Identity* is an instance of self-supervised learning, it can generate its own labeled training data without the user’s involvement. Candidate LTRs outputted by the detector module comprise a database that will be given to *Identity*. We train two instances of *Identity*. The first is trained with a focus on sequence pairs of 80–100% identity scores, and the second is trained with a focus on 60–100% identity scores. We refer to the first *Identity* instance as the standard *Identity* and the second one as the recent *Identity* because it is utilized for locating recently nested retrotransposons. Regarding software integration, *Identity* is accessed through an API and is *not* called externally.

### Matcher

An LTR-retrotransposon is defined by two matched LTRs with an internal part inbetween. This module attempts matching LTRs. Its input is a list of candidate LTRs assembled from the forward stretches and the backward stretches. It outputs a list of LTR-retrotransposon candidates, solo LTR candidates, and complex regions. A solo LTR is defined as a single unmatched LTR. A complex region is characterized by the presence of multiple same-family candidate LTRs. The matcher module follows two steps. It first builds a directed-weighted graph. Using the information from this graph, it matches LTRs.

### Building a Directed-Weighted Graph

*Look4LTRs* utilizes a directed-weighted graph for matching LTR candidates. A graph consists of nodes, which are connected by edges. An edge is directed and has a weight. We add every LTR candidate to the graph as a node. We distinguish between two types of nodes; forward nodes are candidate LTRs from the forward stretches (forward candidates) and backward nodes are candidate LTRs from the backward stretches (backward candidates). An edge between two nodes, i.e., two LTRs, is added when a forward candidate LTR points to a backward candidate and vice versa. Note that no edges are added between two forward candidates or two backward candidates. An edge is assigned a weight representing how similar two candidate LTRs are. Such a similarity is calculated as the ratio of kmers that have copies in the other candidate to the total number of k-mers.

To process recently nested elements, including complete and solo elements, we connect overlapping nodes representing forward and backward candidates. Suppose we have three LTRs nearby each other. The first LTR has a forward node. The third LTR has a backward node. The second node has a forward node (pointing to the third LTR) and a backward node (pointing to the first LTR). Therefore, if a forward and backward node overlap with each other, two edges are added to connect them. We call these edges vertical connections. However, weights associated with vertical connections are not assigned because these connections are solely utilized for connecting LTRs that are nearby each other but are not directly connected.

### Matching LTRs

Our graph consists of a set of connected components. A connected component is a set of connected — directly or indirectly — nodes that are unreachable from any other nodes in the graph. A complete LTR-retrotransposon (with a possible solo LTR nearby) or a group of recently nested elements is represented by a connected component. Figure 4 shows examples of different components. In the simplest case of a single LTR-retrotransposon, a forward node is connected to a backward node, i.e., a 5’ LTR to 3’ LTR.

**Figure 4.**
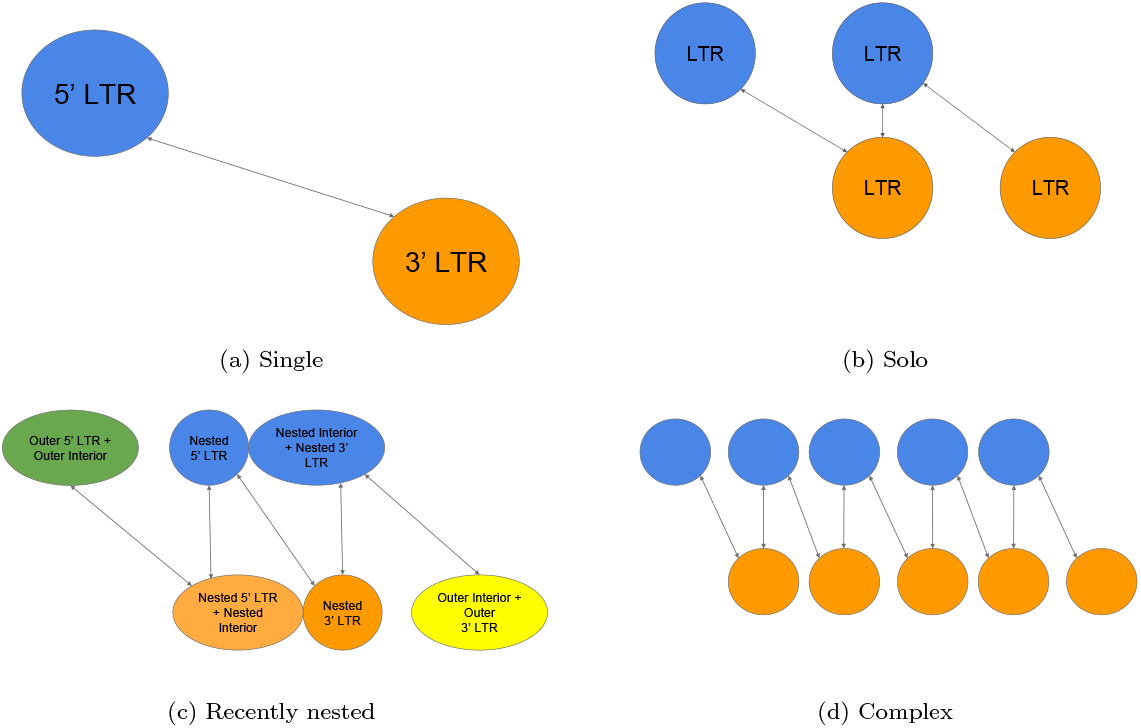
Connected components examples of LTR-retrotransposons. These connected components represent perfect-world scenarios; for simplicity, each edge represents two edges pointing in and out and the weights are not shown. For each example, there are two rows. The top rows contain forward nodes (representing LTR candidates found in the forward stretches) that point forward to a match further in the genome. The bottom rows contain backward nodes (representing LTR candidates found in the backward stretches) that point backwards. When nodes from the forward and backwards nodes overlap, they represent the same LTR. (a) A single LTR-retrotransposon with the 5’ LTR pointing to the 3’ LTR and vice versa. (b) An LTR-retrotransposon with a solo LTR. This connected component may represent one of three scenarios depending on which node represents the solo element, which can be the leftmost node, the rightmost node, or the middle node. (c) A recently nested LTR-retrotransposon. Note that the outermost LTRs merged with the internal part of the outer retrotransposon. Two nodes from the nested LTRs merged with the internal part of the nested retrotransposon. However, two nodes from the nested LTRs are not merged and are distinguishable. (d) A complex graph case where there are many LTRs right by each other.

The merger and the detector modules may wrongfully merge stretches from different repetitive elements, causing an issue we call hyper-extension. When matching nodes together, hyper-extensions can be found by looking at the weights of the connections. If the weight of a connection is low in one direction but high in the other, it may be a sign of this issue. As a remedy, we trim the node from which the edge with the lower weight comes to better match the other node. Then the weights of the connections are recalculated. Figure 5 shows an example of the hyper-extension issue and how it is resolved.

**Figure 5.**
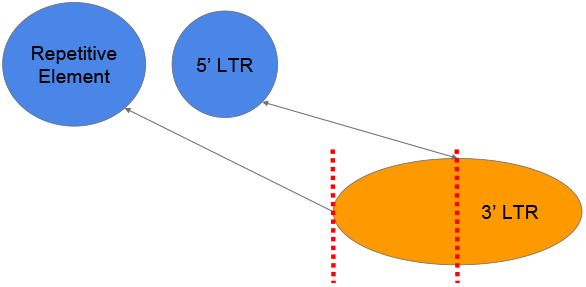
An example of hyper-extension. The 3’ LTR is incorrectly merged with another upstream repetitive element. To remedy this issue, we cut off the region marked with the dotted red lines when matching with the 5’ LTR.

For each subcomponent, we perform case analysis to determine which pair of nodes form an LTR-retrotransposon. We have five cases to consider: (i) the single case, (ii) the solo LTR case, (iii) the recently nested case, (iv) the complex case, and (v) the deeply nested case.

### Single case

In this analysis, we search for single, complete LTR-retrotransposons. First, we look for a forward node that connects to a backward node and vice versa. When found, we check if both weights of the two connections are greater than or equal to 0.27, a value determined by the 95^*th*^ percentile of our training genomes’ ground truth. Then we check for repetitiveness of an internal part using *Red*. We calculate the ratio of non-zero *Red* scores in an internal part to its size. If this value is at least 0.33, determined by the 98^*th*^ percentile on our training genomes’ ground truth, this region is considered a valid internal part. This check ensures that an LTR-retrotransposon candidate follows the general trend for repetitive elements. Because there may be multiple LTRs of the same family nearby, we repeat this process with different combinations of forward and backward nodes that are directly connected, provided they meet the above requirements. Rather than choosing one candidate at this stage, we instead keep multiple pairs and allow the filter module, described later, to filter out false positives.

### Solo case

Here, we search for a solo LTR on either side of a single LTR-retrotransposon or nested within it. Similar to the single case, we check for a minimum weight between the connections of forward and backward nodes. We search for three LTRs where the first is connected to the second and the second is connected to the third. Generally, an internal part of a complete LTR-retrotransposon is expected to be repetitive. Thus, we expect a non-repetitive region to separate a complete LTR-retrotransposon from a solo LTR. *Red* scores are utilized as stated previously to confirm whether a region between LTRs is repetitive. This case is resolved completely if one of the in-between regions is confirmed to be repetitive and the other is not. Otherwise, if all regions are repetitive, it is potentially a nested solo LTR, i.e., the solo LTR is inserted into a complete element of the same family. However, even if all regions are confirmed to be repetitive, it is not certain that the solo LTR is nested. For example, one of the regions could be repetitive because of another inserted unknown repetitive element. In the case where the two in-between regions are repetitive, we output three LTR-retrotransposon candidates, in which the solo LTR can be the first, the second, or the third LTR.

### Recently nested case

In this case, we search for LTR-retrotransposons that are nested within another LTR-retrotransposon of the same family. The boundaries of the outer LTRs are merged with the internal part of the outer element, complicating this analysis. However, the boundaries of the nested LTRs remain intact. The single analysis case can be applied to finding the nested LTR-retrotransposon. Using the nested LTRs, we approximate the boundaries of the outer LTRs using information available in a connected component. At this point, the outermost two LTRs are matched and the innermost two LTRs are matched, resulting in two LTR-retrotransposon candidates. After that, we check if the internal parts of the two LTR-retrotransposon candidates (not including the nested region of the outer LTR-retrotransposon) are repetitive. Because these candidates are supposed to come from the same family, their internal parts should be similar, but not very similar due to expected divergence and other possibly nested elements. For this reason, we diverge from the 80-80-80 rule by lowering the generally used 80% identity score minimum to 60%. We calculate the identity score between the internal parts of the nested and outer retrotransposons using the recent *Identity* model. If the identity score is greater than 60%, we consider these two candidates to be recently nested.

### Complex case

Here, we analyze connected components that have many nearby, same-family elements inserted sequentially. The internal parts of such elements should be similar to each other. To confirm, the regions between nodes (possible internal parts) are gathered and compared to each-other (all vs. all) using the *Identity* standard model. If at least two regions have 80% identity score with each other, we consider the connected component to have sequential LTR-retrotransposons. In essence, this case reports that there are at least two sequentially inserted LTR-retrotransposons. We note that there is no further confirmation of the LTR-retrotransposons for this case due to its complexity.

### Deeply nested case

We previously discussed the recently nested case which handles up to one level of nesting. We designed a deeply nested case to decompose a region consisting of a recently nested element with many levels of nesting. In this case, we apply a recursive process to discover deep nests, i.e., a nested LTR-retrotransposon within another nested LTR-retrotransposon. To begin, we collect a recently nested LTR-retrotransposon found by the recently nested case. We then remove the innermost element from the sequence. After that, all modules starting from the scorer up to the matcher are applied to the newly constructed sequence. This simplifies the deeply nested case to a recursive application of the recently nested case. This process continues until no more recently nested elements are found.

We have discussed how we matched LTRs to form LTR-retrotransposon candidates. We now discuss an intermittent step before the filter module for extending the boundaries of LTRs.

### Extending

The LTRs of an LTR-retrotransposon candidate may not be exact matches, especially length wise. We take two steps to alleviate this issue: (i) k-mer extension and (ii) missing-region extension. In k-mer extension, the 3’ LTR is extended by *k−* 1 bp because the last score is the score of the last k-mer — not the score of the last nucleotide. An LTR may be shorter than its paired LTR in an LTR-retrotransposon candidate. Although this can happen in nature due to insertions and deletions, we attempt to extend the short LTR to account for possible failed merging of stretches in the merger and detector modules. In the missing-region extension, we extend the shorter LTR forward and backward to match the length of the longer LTR. We use *Identity* (the standard) to confirm if an extension results in a better identity score between paired LTRs and keep it if it does.

We discussed how we extend the boundaries of two paired LTRs. In the next stage, candidates are sent to the filter module to drop candidates that fail to meet LTR-retrotransposon’s signature.

### Filter

This module removes LTR-retrotransposon candidates that fail to meet certain criteria such as signature features. The inputs to the filter are LTR-retrotransposon candidates and solo LTR candidates. The outputs are the filtered LTR-retrotransposons and solo LTRs. We check for the following criteria: (i) LTRs and internal parts are of the proper length, (ii) matched LTRs have comparable length and similarity, (iii) LTR-retrotransposon candidates have polypurine tracts (PPT) within the internal parts, and (iv) LTRs are not Miniature Inverted-repeat Transposable Elements (MITEs). A solo LTR is removed at the end if there are no same-family (according to graph information) LTR-retrotransposon candidates that survived.

### Length

The lengths of an LTR-retrotransposon’s components are known to generally be within a certain range. We check if the lengths of the LTRs are at least 2,00 bp and less than 7,000 bp. The internal parts must be at least 2,00 bp long.

### Similarity

The LTRs of an LTR-retrotransposon should have similar sizes and sequences. We check if the length coverage ratio (smaller length to longer length) of two LTRs of an element is at least 0.8. If it is not, we use the Smith-Waterman algorithm to align the two LTRs. If the length of the alignment found is at least 80 bp long and the similarity is at least 80%, we keep the element, otherwise the candidate is removed.

### Polypurine tract

A signature feature of an LTR-retrotransposon is the PPT, which is a sequence of purines (A and G nucleotides), upstream from the 3’ LTR. We take the last 400 bp of an internal part upstream from the 3’ LTR and convert every G to A. Then, we create a sequence of all A nucleotides with a size of 100. Next, the 100-bp-long sequence is aligned to the 400-bp-long sequence using the SmithWaterman algorithm. If the alignment length is greater than 12 and there are more A’s than G’s in the region (a signature of the PPT), the PPT is confirmed. If we do not find a PPT this way, we assume that the orientation is potentially reversed and search downstream using the same process but with T’s and C’s instead of A’s and G’s. If a PPT is not found, the candidate is removed.

### MITE

Here we check for MITEs which are small transposable elements that can be mistaken for LTRs. Further, they are plentiful in plants. Thus, same-family MITEs can be near each other, potentially resulting in the previous modules matching these elements together as LTR candidates into an LTR-retrotransposon candidate. This necessitates a filter to locate and remove them. We check both LTR candidates of the same retrotransposon independently and drop the entire element only if both LTR candidates are found to be potential MITEs. The first and last 30 bp of an LTR candidate are taken and aligned against each other to check if an LTR candidate has the signature feature of a MITE — terminal inverted repeats. If the alignment length found is greater than 15 and the similarity between the two sequences is greater than 85%, we consider it a MITE. We chose the size of 15 because other tools that specialize in MITE detecting using a similar metric of 10 [39, 40]. We took this parameter and increased it by 5 to be stricter.

### Overlapping outputs

After these filters, we consider every LTR-retrotransposon to be a true LTR-retrotransposon. This point should be made clear again; in the matcher module, an LTR may be paired with multiple other LTRs to form LTR-retrotransposons if they meet the criteria. The filter module drops any LTR-retrotransposon that fails to meet the standard features. However, it is possible for multiple LTR-retrotransposons with the same LTR (5’or 3’) to pass every criterion. In this case, all of them are reported.

### Postprocess

LTR-retrotransposon are further sharpened using target site duplications (TSD). We search 20 bp upstream and downstream an LTR-retrotransposon for a duplication and search for the longest common substring between the two regions. If the substring’s length is at least 4, we conclude that we have found a TSD, otherwise it is not reported. We extend the boundaries of the LTR-retrotransposon to meet the TSD if they are not already touching.

### Evaluation

Using our ground truth, we evaluated *Look4LTRs* alongside the following related tools: LTR Finder [41], LTRharvest [27], and LtrDetector [29]. Recall that our ground truth is likely incomplete with possibly inaccurate boundaries. Therefore, the definitions of true positives and false positives needed to be modified to account for these two issues. A true positive is a predicted element that has an 80% reciprocal overlap with an LTR-retrotransposon in the ground truth. For false positives, we followed the method described in previous studies [29, 39]. First, we locate all repetitive elements as reported by RepeatMasker and drop simple repeats and low-complexity regions as well as LTR elements — as described by RepeatMasker. This is our false positive dataset. A false positive is then defined as a predicted element whose two LTRs are overlapping (80% reciprocal) with two elements of the same family from the false positive dataset.

To help determine these true positives and false positives, we utilized BEDTools [42].

The four tools are evaluated by recall, precision, and F1 score. Recall is the percentage of true positives found by a tool. Precision is the percentage of true positives in all confirmed positive predictions — true or false. F1 score is the harmonic mean of recall and precision. Additionally, we report the peak memory usage and the time required to run a tool. The evaluated tools were ran off-the-shelf, utilizing multi-core capabilities only if they came with it because most biologists would apply a tool without any additional modifications. LTRharvest and LTR Finder do not come with multi-core capabilities, whereas *Look4LTRs* and LtrDetector can utilize multiple cores. We used an x86 64 Red Hat Enterprise Linux Server version 7.9 (Maipo) machine to run the tools, utilizing 24 cores if the tool allowed for multiprocessing.

## Results & Discussion

We evaluate *Look4LTRs* alongside these three related tools: LTR Finder [41], LTRharvest [27], and LtrDetector [29]. *Look4LTRs* was evaluated on the four training genomes and the four testing genomes. The performance due to the cross-species feature of *Look4LTRs* was compared to the other tools’ performances on five genomes from the rice family.

### Training & Testing Evaluation

Our training genomes consist of *Arabidopsis thaliana, Oryza sativa japonica, Glycine max*, and *Sorghum bicolor*. The testing genomes consist of *Zea mays, Solanum lycopersicum, Solanum tuberosum*, and *Theobroma cacao*. Table 1 shows the results of the tools on the training and testing genomes. Due to time constraints, we did not run LTR Finder on *Zea mays*. Previous experience with LTR Finder shows that *Zea mays* would take many days of non-interrupted running to complete. We will now discuss the results on the training and testing genomes.

**Table 1.**
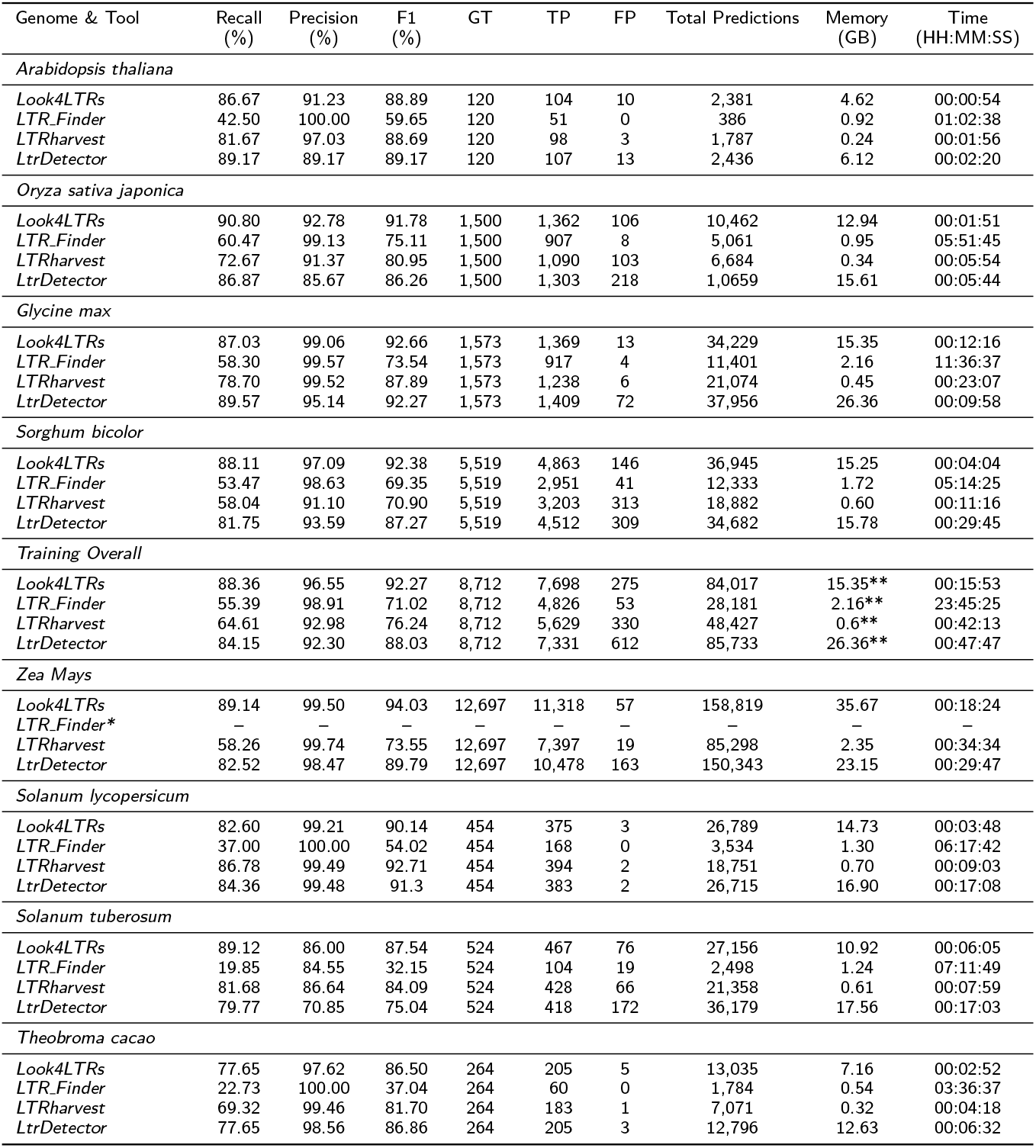
Results of *Look4LTRs*, LTR Finder, LTRharvest, and LtrDetector on the training and testing genomes. GT is the number of elements in the ground truth, TP is the number of true positives, FP is the number of false positives, Total Predictions is the number of predictions made by a tool, memory is the peak memory usage of a tool, and time is the total time taken by a tool to run. *We did not run LTR Finder on *Zea mays* due to the time it would take. **For the overall peak memories, we take the maximum peak memory usage from each tool over the training genomes. Note that LTRharvest and LTR Finder were run as provided by their authors — without parallelization — because a biologist would apply them this way without any modifications.

| Genome & Tool | Recall (%) | Precision (%) | F1 (%) | GT | TP | FP | Total Predictions | Memory (GB) | Time (HH:MM:SS) |
| --- | --- | --- | --- | --- | --- | --- | --- | --- | --- |
| *Arabidopsis thaliana* |  |  |  |  |  |  |  |  |  |
| *Look4LTRs* | 86.67 | 91.23 | 88.89 | 120 | 104 | 10 | 2,381 | 4.62 | 00:00:54 |
| *LTR.Finder* | 42.50 | 100.00 | 59.65 | 120 | 51 | 0 | 386 | 0.92 | 01:02:38 |
| *LTRharvest* | 81.67 | 97.03 | 88.69 | 120 | 98 | 3 | 1,787 | 0.24 | 00:01:56 |
| *LtrDetector* | 89.17 | 89.17 | 89.17 | 120 | 107 | 13 | 2,436 | 6.12 | 00:02:20 |
| *Oryza sativa japonica* |  |  |  |  |  |  |  |  |  |
| *Look4LTRs* | 90.80 | 92.78 | 91.78 | 1,500 | 1,362 | 106 | 10,462 | 12.94 | 00:01:51 |
| *LTR.Finder* | 60.47 | 99.13 | 75.11 | 1,500 | 907 | 8 | 5,061 | 0.95 | 05:51:45 |
| *LTRharvest* | 72.67 | 91.37 | 80.95 | 1,500 | 1,090 | 103 | 6,684 | 0.34 | 00:05:54 |
| *LtrDetector* | 86.87 | 85.67 | 86.26 | 1,500 | 1,303 | 218 | 1,0659 | 15.61 | 00:05:44 |
| *Glycine max* |  |  |  |  |  |  |  |  |  |
| *Look4LTRs* | 87.03 | 99.06 | 92.66 | 1,573 | 1,369 | 13 | 34,229 | 15.35 | 00:12:16 |
| *LTR.Finder* | 58.30 | 99.57 | 73.54 | 1,573 | 917 | 4 | 11,401 | 2.16 | 11:36:37 |
| *LTRharvest* | 78.70 | 99.52 | 87.89 | 1,573 | 1,238 | 6 | 21,074 | 0.45 | 00:23:07 |
| *LtrDetector* | 89.57 | 95.14 | 92.27 | 1,573 | 1,409 | 72 | 37,956 | 26.36 | 00:09:58 |
| *Sorghum bicolor* |  |  |  |  |  |  |  |  |  |
| *Look4LTRs* | 88.11 | 97.09 | 92.38 | 5,519 | 4,863 | 146 | 36,945 | 15.25 | 00:04:04 |
| *LTR.Finder* | 53.47 | 98.63 | 69.35 | 5,519 | 2,951 | 41 | 12,333 | 1.72 | 05:14:25 |
| *LTRharvest* | 58.04 | 91.10 | 70.90 | 5,519 | 3,203 | 313 | 18,882 | 0.60 | 00:11:16 |
| *LtrDetector* | 81.75 | 93.59 | 87.27 | 5,519 | 4,512 | 309 | 34,682 | 15.78 | 00:29:45 |
| *Training Overall* |  |  |  |  |  |  |  |  |  |
| *Look4LTRs* | 88.36 | 96.55 | 92.27 | 8,712 | 7,698 | 275 | 84,017 | 15.35** | 00:15:53 |
| *LTR.Finder* | 55.39 | 98.91 | 71.02 | 8,712 | 4,826 | 53 | 28,181 | 2.16** | 23:45:25 |
| *LTRharvest* | 64.61 | 92.98 | 76.24 | 8,712 | 5,629 | 330 | 48,427 | 0.6** | 00:42:13 |
| *LtrDetector* | 84.15 | 92.30 | 88.03 | 8,712 | 7,331 | 612 | 85,733 | 26.36** | 00:47:47 |
| *Zea Mays* |  |  |  |  |  |  |  |  |  |
| *Look4LTRs* | 89.14 | 99.50 | 94.03 | 12,697 | 11,318 | 57 | 158,819 | 35.67 | 00:18:24 |
| *LTR.Finder\** | — | — | — | — | — | — | — | — | — |
| *LTRharvest* | 58.26 | 99.74 | 73.55 | 12,697 | 7,397 | 19 | 85,298 | 2.35 | 00:34:34 |
| *LtrDetector* | 82.52 | 98.47 | 89.79 | 12,697 | 10,478 | 163 | 150,343 | 23.15 | 00:29:47 |
| *Solanum lycopersicum* |  |  |  |  |  |  |  |  |  |
| *Look4LTRs* | 82.60 | 99.21 | 90.14 | 454 | 375 | 3 | 26,789 | 14.73 | 00:03:48 |
| *LTR.Finder* | 37.00 | 100.00 | 54.02 | 454 | 168 | 0 | 3,534 | 1.30 | 06:17:42 |
| *LTRharvest* | 86.78 | 99.49 | 92.71 | 454 | 394 | 2 | 18,751 | 0.70 | 00:09:03 |
| *LtrDetector* | 84.36 | 99.48 | 91.3 | 454 | 383 | 2 | 26,715 | 16.90 | 00:17:08 |
| *Solanum tuberosum* |  |  |  |  |  |  |  |  |  |
| *Look4LTRs* | 89.12 | 86.00 | 87.54 | 524 | 467 | 76 | 27,156 | 10.92 | 00:06:05 |
| *LTR.Finder* | 19.85 | 84.55 | 32.15 | 524 | 104 | 19 | 2,498 | 1.24 | 07:11:49 |
| *LTRharvest* | 81.68 | 86.64 | 84.09 | 524 | 428 | 66 | 21,358 | 0.61 | 00:07:59 |
| *LtrDetector* | 79.77 | 70.85 | 75.04 | 524 | 418 | 172 | 36,179 | 17.56 | 00:17:03 |
| *Theobroma cacao* |  |  |  |  |  |  |  |  |  |
| *Look4LTRs* | 77.65 | 97.62 | 86.50 | 264 | 205 | 5 | 13,035 | 7.16 | 00:02:52 |
| *LTR.Finder* | 22.73 | 100.00 | 37.04 | 264 | 60 | 0 | 1,784 | 0.54 | 03:36:37 |
| *LTRharvest* | 69.32 | 99.46 | 81.70 | 264 | 183 | 1 | 7,071 | 0.32 | 00:04:18 |
| *LtrDetector* | 77.65 | 98.56 | 86.86 | 264 | 205 | 3 | 12,796 | 12.63 | 00:06:32 |

**Table 2.**
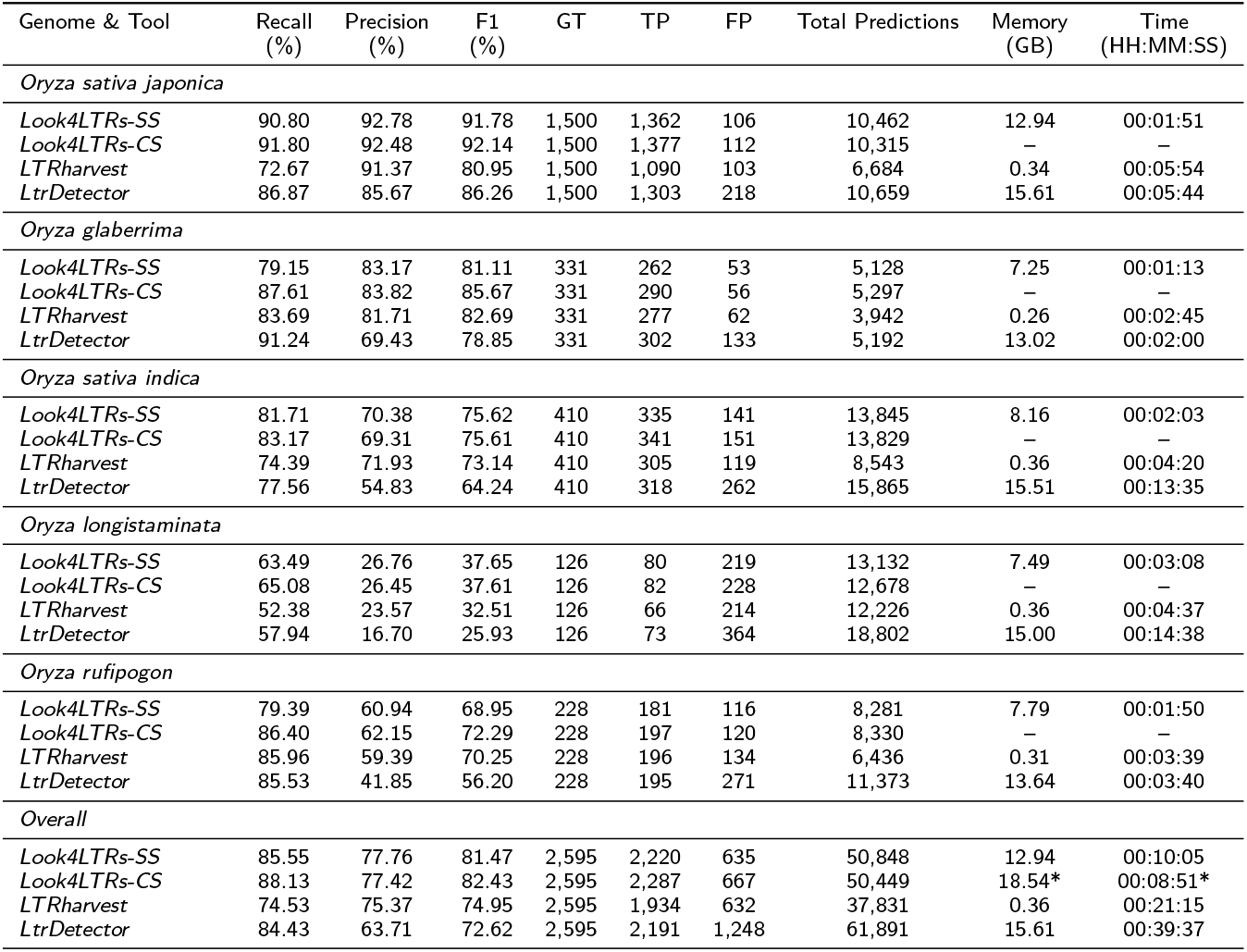
Cross-species results of *Look4LTRs* compared to other tools. The tools used are *Look4LTRs-SS* (Single-Species), *Look4LTRs-CS* (Cross-Species), LTRharvest, and LtrDetector on the following five rice species: (i) *Oryza sativa japonica*, (ii) *Oryza glaberrima*, (iii) *Oryza sativa indica*, (iv) *Oryza longistaminata*, and (v) *Oryza rufipogon. Look4LTRs-SS* is applied to each species separately, i.e., it only trains and predicts on one given genome at a time. *Look4LTRs-CS* is applied to the five species together, i.e., it learns the repeat content of the five species and predicts on all of them. *The memory and time for *Look4LTRs-CS* is the same for every genome and the Overall because *Look4LTRs-CS* was run on all genomes at once. Thus, they all come from the same exact run.

| Genome & Tool | Recall (%) | Precision (%) | F1 (%) | GT | TP | FP | Total Predictions | Memory (GB) | Time (HH:MM:SS) |
| --- | --- | --- | --- | --- | --- | --- | --- | --- | --- |
| *Oryza sativa japonica* |  |  |  |  |  |  |  |  |  |
| *Look4LTRs*-SS | 90.80 | 92.78 | 91.78 | 1,500 | 1,362 | 106 | 10,462 | 12.94 | 00:01:51 |
| *Look4LTRs*-CS | 91.80 | 92.48 | 92.14 | 1,500 | 1,377 | 112 | 10,315 | — | — |
| LTRharvest | 72.67 | 91.37 | 80.95 | 1,500 | 1,090 | 103 | 6,684 | 0.34 | 00:05:54 |
| LtrDetector | 86.87 | 85.67 | 86.26 | 1,500 | 1,303 | 218 | 10,659 | 15.61 | 00:05:44 |
| *Oryza glaberrima* |  |  |  |  |  |  |  |  |  |
| *Look4LTRs*-SS | 79.15 | 83.17 | 81.11 | 331 | 262 | 53 | 5,128 | 7.25 | 00:01:13 |
| *Look4LTRs*-CS | 87.61 | 83.82 | 85.67 | 331 | 290 | 56 | 5,297 | — | — |
| LTRharvest | 83.69 | 81.71 | 82.69 | 331 | 277 | 62 | 3,942 | 0.26 | 00:02:45 |
| LtrDetector | 91.24 | 69.43 | 78.85 | 331 | 302 | 133 | 5,192 | 13.02 | 00:02:00 |
| *Oryza sativa indica* |  |  |  |  |  |  |  |  |  |
| *Look4LTRs*-SS | 81.71 | 70.38 | 75.62 | 410 | 335 | 141 | 13,845 | 8.16 | 00:02:03 |
| *Look4LTRs*-CS | 83.17 | 69.31 | 75.61 | 410 | 341 | 151 | 13,829 | — | — |
| LTRharvest | 74.39 | 71.93 | 73.14 | 410 | 305 | 119 | 8,543 | 0.36 | 00:04:20 |
| LtrDetector | 77.56 | 54.83 | 64.24 | 410 | 318 | 262 | 15,865 | 15.51 | 00:13:35 |
| *Oryza longistaminata* |  |  |  |  |  |  |  |  |  |
| *Look4LTRs*-SS | 63.49 | 26.76 | 37.65 | 126 | 80 | 219 | 13,132 | 7.49 | 00:03:08 |
| *Look4LTRs*-CS | 65.08 | 26.45 | 37.61 | 126 | 82 | 228 | 12,678 | — | — |
| LTRharvest | 52.38 | 23.57 | 32.51 | 126 | 66 | 214 | 12,226 | 0.36 | 00:04:37 |
| LtrDetector | 57.94 | 16.70 | 25.93 | 126 | 73 | 364 | 18,802 | 15.00 | 00:14:38 |
| *Oryza rufipogon* |  |  |  |  |  |  |  |  |  |
| *Look4LTRs*-SS | 79.39 | 60.94 | 68.95 | 228 | 181 | 116 | 8,281 | 7.79 | 00:01:50 |
| *Look4LTRs*-CS | 86.40 | 62.15 | 72.29 | 228 | 197 | 120 | 8,330 | — | — |
| LTRharvest | 85.96 | 59.39 | 70.25 | 228 | 196 | 134 | 6,436 | 0.31 | 00:03:39 |
| LtrDetector | 85.53 | 41.85 | 56.20 | 228 | 195 | 271 | 11,373 | 13.64 | 00:03:40 |
| *Overall* |  |  |  |  |  |  |  |  |  |
| *Look4LTRs*-SS | 85.55 | 77.76 | 81.47 | 2,595 | 2,220 | 635 | 50,848 | 12.94 | 00:10:05 |
| *Look4LTRs*-CS | 88.13 | 77.42 | 82.43 | 2,595 | 2,287 | 667 | 50,449 | 18.54* | 00:08:51* |
| LTRharvest | 74.53 | 75.37 | 74.95 | 2,595 | 1,934 | 632 | 37,831 | 0.36 | 00:21:15 |
| LtrDetector | 84.43 | 63.71 | 72.62 | 2,595 | 2,191 | 1,248 | 61,891 | 15.61 | 00:39:37 |

### Recall

On the training genomes, *Look4LTRs* was the best in terms of recall overall, followed by LtrDetector, LTRharvest, and then LTR Finder. On the testing genomes, *Look4LTRs* came first on three genomes and third on one genome (*Solanum lycopersicum*). However, it was notably in first place on the largest genome of the four (*Zea mays*) showing an improvement from the second-best tool (LtrDetector) by 8%. On *Theobroma cacao, Look4LTRs* tied for first place with LtrDetector. LtrDetector and LTRharvest varied between first, second, and third place on the testing genomes. LTR Finder was consistently in last place in terms of recall. Thus, *Look4LTRs* is highly capable of finding LTR-retrotransposons; *Look4LTRs* is comparable to the other tools and surpasses them in many cases.

### Precision

Overall, LTR Finder was the best in terms of precision on the training genomes, followed by *Look4LTRs*, LTRharvest, and LtrDetector (comparable to LTRharvest). All tools achieved high overall precision scores (92–99%) on the training genomes. Next, we discuss precision scores on the testing genomes. On *Zea mays*, LTRharvest was in first place; nonetheless, *Look4LTRs* (in second place), and LtrDetector (third place) were very comparable. The results for *Solanum lycopersicum* show a similar trend with LTR Finder in first place (100%) but the other tools were very comparable — LTRharvest at 99.49%, LtrDetector at 99.48%, and *Look4LTRs* at 99.21%. For *Solanum tuberosum*, all tools suffered in precision; however, all but LtrDetector (70.85%) were very comparable (84.55–86.64%). Finally, on *Theobroma cacao, Look4LTRs* achieved 97.62%; however, *Look4LTRs* came last because all other tools were highly precise. *Look4LTRs* is very comparable to the other tools in terms of precision. On the genomes where it fell behind the others, the difference was minimal.

### F1 score

*Look4LTRs* was the best in terms of F1 score, followed by LtrDetector, LTRharvest, and then LTR Finder, collectively on the training genomes. With respect to the testing genomes, *Look4LTRs* came first (94.03%) on *Zea mays*, followed by LtrDetector (89.79%) and LTRharvest (73.55%). For *Solanum lycopersicum*, LTRharvest came first at 92.71%, but was comparable to LtrDetector (91.3%) and *Look4LTRs* (90.14%); LTR Finder came last at 54.02%. *Look4LTRs* was in first place at 87.54% on *Solanum tuberosum*, closely followed by LTRharvest at 84.09%; LtrDetector was in third place at 75.05% and LTR Finder was in last place at 32.15%. For *Theobroma cacao*, LtrDetector was in first place at 86.86%, followed closely by *Look4LTRs* at 86.5%, then followed by LTRharvest at 81.7% and LTR Finder at 37.04%. From these results, *Look4LTRs* offers a good balance of recall and precision evident by achieving the highest overall F1 score on the training genomes and either the highest F1 score or comparable scores to those of the best performing tools on the testing genomes.

### Number of predictions

Overall, on the training genomes, LTR Finder made the least number of predictions at 28,181, followed by LTRharvest at 48,427. *Look4LTRs* produced 84,017 predictions and LtrDetector produced a comparable amount of 85,733 predictions. This trend continued for every genome — training or testing — where LTR Finder makes the least number of predictions, followed by LTRharvest. *Look4LTRs* and LtrDetector predict a comparable number of predictions and vary between third and fourth place on the testing genomes. *Look4LTRs* produces less predictions than LtrDetector while retaining a high recall and F1 score.

### Peak memory usage

The lowest peak memory overall on the training genomes was LTRharvest at 0.6 GigaBytes (GBs), followed by LTR Finder at 1.72 GBs, Look4LTRs at 15.35 GBs, and LtrDetector at 26.36 GBs. For the results on the training genomes, the same trend continues except for on *Zea mays* where *Look4LTRs* takes 35.67 GBs and LtrDetector takes 23.15 GBs. *Look4LTRs* takes less memory than LtrDetector on most genomes. Although it was not the lowest in memory consumption, many modern computational machines can handle the memory requirements of *Look4LTRs*.

### Time

*Look4LTRs* was the quickest on the training genomes overall at approximately 16 minutes, followed by LTRharvest at 42 minutes, LtrDetector at 47 minutes, and LTR Finder at 23 hours. In regards to the testing genomes, *Look4LTRs* was the quickest on *Zea mays* at 18 minutes, followed by LtrDetector at nearly 30 minutes and LTRharvest at 34 minutes. For the rest of the testing genomes, *Look4LTRs* was also the quickest followed by LTRharvest, LtrDetector, and then LTR Finder. *Look4LTRs* was quick in comparison to the other tools. We note, however, that there is a trade-off in speed to memory usage. The more the cores, the more the memory requirements; if time is not an issue, then *Look4LTRs* can be run on a single or few cores to reduce memory usage at the cost of *Look4LTRs*’ quick speed.

### Cross-Species Evaluation

A crucial feature of *Look4LTRs* is its suitability to cross-species studies. We evaluated *Look4LTRs* on five species from the rice (*Oryza*) family. These five species are: (i) *Oryza glaberrima*, (ii) *Oryza sativa indica*, (iii) *Oryza longistaminata*, (iv) *Oryza rufipogon*, and (v) *Oryza sativa japonica*. The ground truths for these species — except *Oryza sativa japonica* and *Oryza sativa indica* — were generated as previously mentioned with one slight modification. As these species are not specifically well annotated in RepeatMasker’s Repbase library, we pass *Oryza* as the species parameter of RepeatMasker. As mentioned before, *Look4LTRs* trained on *Oryza sativa japonica*; however, we did not train the cross-species component of *Look4LTRs* on *Oryza sativa japonica*. We also evaluated LTRharvest and LtrDetector on these species for comparison.

We use two different versions of *Look4LTRs* to evaluate. The first version of *Look4LTRs*, which we refer to as *Look4LTRs-SS* (Single Species), is applied to each species separately, i.e., it only trains and predicts on one given genome at a time. The second version of *Look4LTRs*, which we refer to as *Look4LTRs-CS* (Cross Species), is applied to the five species together, i.e., it trains on the five species and predicts on all of them. The purpose of these two applications is to study the added benefit of using information from related-species for finding LTR-retrotransposons. Table 2 displays the results of the tools on the mentioned rice species.

Overall, *Look4LTRs-CS* has the highest recall and F1 score. *Look4LTRs-CS* shows an improvement over *Look4LTRs-SS*. The cross-species aspect of *Look4LTRs* improved the overall recall (88.55% vs. 85.13%). The precision of the cross-species version was very comparable to the single-species version (77.42% vs. 77.76%). For the overall F1 score, *Look4LTRs-CS* was at 82.43% in comparison to *Look4LTRs-SS* at 81.47%. This slight improvement was due to the improvement in recall. Additionally, there was a speed-up advantage for using the cross-species version when running all of the genomes; *Look4LTRs-CS* took approximately 9 minutes, whereas *Look4LTRs-SS* took 10 minutes, LTRharvest took 21 minutes, and LtrDetector took nearly 40 minutes. These results demonstrate that information from related species utilized in *Look4LTRs-CS* improved the discovery of LTR-retrotransposons.

### Coverage estimation of LTR-retrotransposons

We estimated the coverage of LTR-retrotransposons in the twelve analyzed genomes and an additional genome of *Hordeum vulgare*. We used the output of *Look4LTRs* for all but the rice genomes, which we instead used the output of *Look4LTRs-CS*. Figure 6 shows these percentages. Interestingly, *Oryza sativa indica*, a subspecies of *Oryza sativa*, has more LTR-retrotransposons than the other subspecies *Oryza sativa japonica* (30.78% vs. 22.14% coverage).

**Figure 6.**
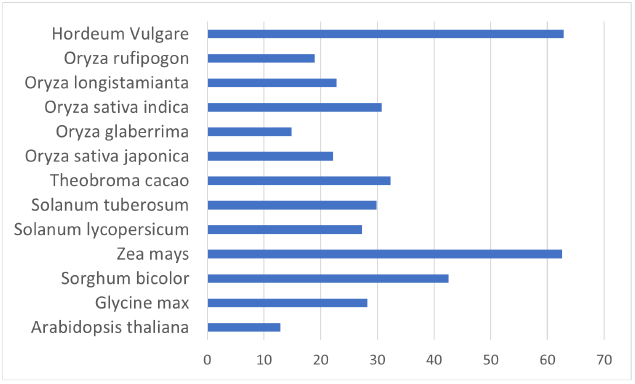
Coverage of LTR-retrotransposons in the analyzed genomes. The coverage is the percentage of the genome that is covered by LTR-retrotransposons. LTR-retrotransposons comprise more than 60% of *Zea mays* and *Hordeum vulgare* genomes.

*Look4LTRs* is able to discover recently nested repeats; we show our estimation of their coverage in the thirteen genomes in Figure 7. We found that *Sorghum bicolor* has the highest content of recently nested LTR-retrotransposons at 0.53%, followed by *Glycine max* at 0.38%, *Theobroma cacao* at 0.36%, *Hordeum vulgare* at 0.21%, and *Zea mays* at 0.08%. On the remaining genomes, *Look4LTRs* was only able to find very few recently nested LTR-retrotransposons. On *Oryza sativa indica* and *Oryza rufipogon, Look4LTRs* was unable to find any recently nested LTR-retrotransposons.

**Figure 7.**
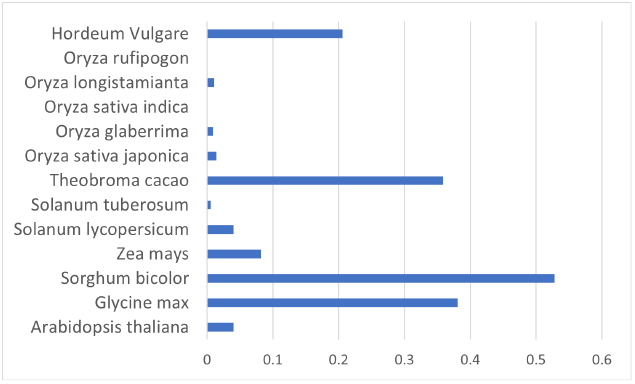
Coverage of recently nested LTR-retrotransposons in the analyzed genomes. The coverage is the percentage of the genome that is covered by recently nested LTR-retrotransposons. Due to the difficulty in detangling recently nested LTR-retrotransposons, these recently nested elements found may be less than the true number of recently nested elements. *Oryza sativa indica* and *Oryza rufipogon*, where no recently nested elements were found by *Look4LTRs*, are a good example of the difficulty in finding recently nested elements. In *Sorghum bicolor*, 0.53% of the genome is covered by recently nested LTR-retrotransposons.

### Expert manual confirmation

A blind manual evaluation was performed on the results of *Look4LTRs* applied to the barley genome. This plant has a haploid genome size of about 5.3 Gb distributed in seven chromosomes with a TE content of about 76% composed in majority of LTR-retrotransposons [43]. Sequences identified as single LTR-retrotransposons and recently nested insertions were investigated. For this, each sequence was compared to the Repbase database and to the NCBI nonredundant protein database using BlastX [38].

Concerning single LTR-retrotransposons, six sequences were assessed. Three of them correspond to known LTR-retrotransposons (BARE1, BARE-2 and Sabrina) described in the barley. Two sequences could correspond to new families not described yet that contain a pol gene and two sequences likely corresponding to LTRs at each extremity, while the last sequence is a false positive not corresponding to any TE. From the two sequences from potentially new families, the first sequence is located in chromosome 2H, starting at position 252,757,149 and ending at position 252,771,106. The second sequence is located in chromosome 3H, starting at position 380,967,486 and ending at position 380,979,130.

In addition, we checked five sequences considered by *Look4LTRs* as recently nested elements. Two of them correspond to false positives. However, the three other sequences correspond to a LTR-retrotransposon inserted into another one (Figure 8). More specifically, two cases correspond to an LTR-retrotransposon (BARE-2 and BARE1) inserted into one LTR of a BARE1 element. The last case represents a BARE1 element inserted into the internal part of another BARE1 element. In the first case in chromosome 1H, the outer element (BARE1) is at position 459,838,459 and ends at position 459,853,480. The nested element (BARE-2) is at position 459,839,301 and ends at 459,847,888. In the second case, the outer element is at position 630,335,812 and ends at 630,353,010. The nested element is at position 630,337,503 and ends at 630,346,445. In the third case, the outer element is at position 564,092,881 and ends at 564,110,748. The nested element is at position 564,095,106 and ends at 564,104,043. This evaluation shows that *Look4LTRs* is able to find recently nested LTR-retrotransposons.

**Figure 8.**
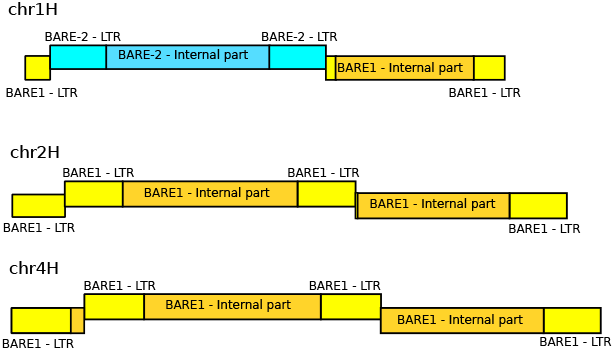
Three cases of recently nested LTR-retrotransposons located in *Hordeum vulgare* on chromosomes 1H, 2H and 4H identified by *Look4LTRs*.

## Conclusion

*Look4LTRs* is a novel tool for LTR-retrotransposon discovery. It processes a whole genome or a group of related genomes simultaneously. *Look4LTRs* usage of the repetitive content of genomes allows for cross-species studies by finding shared elements between closely related species. According to our evaluations, the usage of repetitive content from related species improves the recall and F1 scores. The repetitive content learned is also used as evidence to enforce that predicted LTR-retrotransposons are repetitive, adhering to the definition of a TE. Further, *Look4LTRs* is able to find recently nested LTR-retrotransposons. Through this feature, we have determined that 0.53% of *Sorghum bicolor* is made of recently nested LTR-retrotransposons.

*Look4LTRs* is nearly alignment free, depending on a kmer matching technique and a graph-based algorithm to find and match LTRs. As a result, *Look4LTRs* has a low runtime, capable of processing the *Zea mays* genome. We are convinced that *Look4LTRs* will be a great addition to the LTR-retrotransposon detection toolset due to the novel features it presents.

## Funding

We were supported by internal funding from the Texas A&M University-Kingsville College of Engineering.

### Abbreviations

TE: Transposable Element
LTR: Long Terminal Repeat
TSD: Target Site Duplication
PPT: Polypurine Tract
bp: base pair

## Availability of data and materials

Tool name: *Look4LTRs*

Github: https://github.com/BioinformaticsToolsmith/Look4LTRs

Operating System: UNIX/LINUX

Programming Language: C++

License: GNU Affero General Public License v3.0 Any restrictions to use by

non-academics: Alternative commercial license is required

## Ethics approval and consent to participate

Not applicable.

## Consent for publication

Not applicable.

## Authors’ contributions

Hani Z. Girgis provided the concept behind *Look4LTRs*, helped developed *Look4LTRs*, evaluated *Look4LTRs*, and aided in writing the manuscript. Anthony B. Garza developed *Look4LTRs*, evaluated *Look4LTRs*, and wrote the manuscript. Lerat Emmanuelle provided biological insight on the parameters and a blind manual expertise on the results of Look4LTRs applied to the barley genome.

## Authors’ information

Anthony B. Garza and Hani Z. Girgis.

Bioinformatics Toolsmith Laboratory,

Department of Electrical Engineering and Computer Science, Texas A&M University-Kingsville, Kingsville, TX, USA

Lerat Emmanuelle.

The Biometrics and Evolutionary Biology Laboratory, University Lyon 1, Villeurbanne, France

